# Codon Usage Bias Analysis of the *WRKY* Gene Family in *Musa acuminata*

**DOI:** 10.1101/2025.01.08.631839

**Authors:** Jiaman Sun, Ji Zhang, Jinzhong Zhang, Brett J. Ferguson, Andrew Chen

**Affiliations:** Guangdong Provincial Key Laboratory of Conservation and Precision Utilization of Characteristic Agricultural Resources in Mountainous Areas, School of Life Science, Jiaying University, Meizhou, Guangdong, 514015, China; Guangxi Academy of Agricultural Sciences, Nanning 530007, China; Integrative Legume Research Group, School of Agriculture and Food Sustainability, The University of Queensland, St. Lucia, QLD 4072, Australia; School of Agriculture and Food Sustainability, The University of Queensland, St Lucia, Brisbane, QLD 4072, Australia

**Keywords:** *Musa acuminata*, ’Guijiao 9’, *WRKY* transcription factors, codon usage bias, natural selection, mutation pressure, Fusarium wilt of banana, *Fusarium oxysporum* f. sp. *cubense*, Tropical Race 4

## Abstract

Codon usage bias (CUB) is a universal phenomenon across all organisms and species, providing valuable insights into codon usage patterns to understand evolutionary mechanisms. The *WRKY* transcription factor family plays a pivotal role in regulating plant responses to various stresses at both physiological and biochemical levels. Bananas (*Musa* spp.), widely consumed as a fruit or staple food, are an important horticulture crop in the tropical and subtropical regions around the world. A total of 151 *WRKY* transcription factors were identified across the genome of the *Musa acuminata* banana cultivar ‘Guijiao 9’, which has been shown to be resistant to the Tropical Race 4 of the Fusarium wilt of banana. The codons of these transcription factors, selected based on their expression from RNA-Seq data, were analysed to investigate the evolutionary patterns of WRKY genes in *Musa acuminata*. The average GC content of *MaWRKY* genes was 55.49%, with a GC3 content of 62.13%, indicating a preference for G/C-ending codons. Among the codons, 27 were identified as high frequency, with 22 ending in G or C. The high effective number of codons (ENC) values (34.53–60.79) and low codon adaptation index (CAI) values (0.143–0.3) suggested weak CUB and that was associated with low gene expression levels. ENC-plot, PR2-plot, and neutrality analysis revealed that both natural selection and mutation pressure contributed to the observed CUB, with natural selection being the dominant factor influencing the codon usage of *MaWRKY* genes in *M. acuminata* ‘Guijiao 9’. Fourteen optimal codons, all ending in G or C, were identified. This analysis provides a theoretical foundation for further understanding the evolutionary mechanisms of *WRKY* genes and identifying key genes involved in resistance to biotic stress in *Musa*. Analysing the evolution of *WRKY* family genes can offer valuable genetic insights into banana growth and development, supporting the phenotypic selection of banana hybrids in breeding programs.

## Introduction

A codon is a triad nucleotide sequence that encodes amino acids on messenger RNA. Codons that encode the same amino acid are referred to as synonymous codons. Over the course of evolution, certain species and specific genes tend to favour particular codons for encoding amino acids, leading to synonymous codon usage bias (Kumar et al., 2016). This bias influences gene expression levels, with stronger codon usage bias generally correlating with higher gene expression (Jiang et al., 2008). Low-frequency codons help maintain codon diversity and contribute to gene expression regulation, playing a critical role in species stability and evolution of each species (Liu et al., 2010). Three primary forces shaping codon usage bias have been identified: mutation pressure, natural selection and random genetic drift (Behura and Severson, 2013). Mutation pressure causes deviations in nucleotide composition, while natural selection restricts codon usage bias to optimise protein production efficiency in highly expressed genes (Guan et al., 2018). Additional factors influencing codon usage bias include nucleic acid composition (Mondal et al., 2016), gene expression level (Hambuch and Parsch, 2005), gene length (Wei et al., 2014), and tRNA abundance (Deng et al., 2020).Analysis of codon usage bias is crucial for understanding the evolutionary mechanisms and codon usage patterns of specific plants. Synonymous codon usage bias differs between monocotyledons and dicotyledons. Monocotyledons tend to favour codons ending in G/C, while dicotyledons prefer A/T endings (Murray et al., 1989). GC content also serves as a distinguishing factor between these two groups. Related species often exhibit similar codon usage biases, making it an important tool for inferring evolutionary relationships among different plant species (Ma et al., 2015).

Codon usage bias reflects the origins, evolution, and mutations of species or genes, providing essential insights for gene function analysis, protein expression and protein structure research (Wu et al., 2015; He et al., 2016). In recent years, studies on codon preference in gene families and individual genes across plants (Liu et al., 2020), animals (Uddin et al., 2020), viruses (Deb et al., 2020; Xin et al., 2020) and other organisms has been increasingly widespread. *WRKY* transcription factor family is one of the largest transcription factor families in plants, playing a key role in physiological and biochemical processes such as hormone synthesis, signal transduction, plant growth and development, metabolism regulation and biological responses to both biotic and abiotic stresses (Govardhana and Satyan, 2020; Li et al., 2020). The widespread use of next-generation sequencing technologies for whole genome and transcriptome sequencing, along with the availability of bioinformatics tools have greatly advanced in-depth studies of WRKY transcription factor family genes and their functional roles in plant’s ability to adapt to environmental challenges (Wani et al., 2021; Goyal et al., 2023; Liu et al., 2023; Deng et al., 2024).Banana (*Musa.* spp) is an important fruit crop, which supplies essential nutrients to millions of people around the world (Drenth and Kema, 2021). Cultivated bananas originated from natural intra-and interspecific hybridisation between two diploid species, *Musa acuminata (*AA genome) and *Musa balbisiana* (BB genome) (Drenth and Kema, 2021). However, due to various biotic and abiotic stresses, global banana yields have significantly declined in recent years (Siamak and Zheng, 2018; Patel et al., 2019).

One of the major diseases threatening the banana production dominated by ‘Cavendish’ from the AAA subgroup is the Fusarium wilt of banana, caused by the *Fusarium oxysporum* f. sp. *cubense* tropical race 4 (*Foc* TR4) (Zorrilla-Fontanesi et al., 2020; Roberts et al., 2024). Current efforts have been focused on containment and deterrence through biosecurity measures; biological control agents; integrated disease management strategies; and the development of resistant banana varieties (Swarupa et al., 2014; Ploetz, 2015; Dita et al., 2018; Pegg et al., 2019). Resistant (R) genes, such as Resistance Gene Analog 2 (RGA2), which was isolated from *Foc TR4*-resistant *Musa acuminata* ssp. *malaccensis*, demonstrated strong resistance in the field when over-expressed in ‘Cavendish’ transgenic lines (Dale et al., 2017). Forward genetic studies have also identified QTLs conferring resistance to *Foc* TR4 and subtropical race 4 (STR4) (Ahmad et al., 2020; Chen et al., 2023a, 2023b). Candidate R genes, specifically pattern recognition receptors, were identified.

Several *WRKY* genes, such as *MaWRKY28, MaWRKY71, MaWRKY40 and MaWRKY22,* have been associated with banana’s response to biotic stress, including *Foc* TR4 (Sun et al., 2019; Li et al., 2012). However, research on the codon usage pattern of WRKY genes in banana has not been reported. In our previous study, *Musa. acuminata* banana ‘Guijiao 9’ exhibited strong resistance to *Foc*-TR4, with lower disease incidence and severity compared to susceptible banana cultivars (Sun et al., 2019). In this study, 151 members of WRKY gene family were identified from *M. acuminata* ‘Guijiao 9’ transcriptome data. Factors influencing codon preference of *MaWRKY* and potential evolutionary models were determined by analysing the CUB of *MaWRKY* genes in ‘Guijiao 9’. These findings contribute to understanding the function of the *WRKY* gene family and provide insights for codon optimisation of *MaWRKY* in the process of biotic stress.

## Results

### Codon composition of *MaWRKY* genes reveals a G/C bias

The codon composition and ENC values of 151 banana *MaWRKY* family members were statistically analysed, with detailed codon bias parameters provided in Additional File 2. The ENC values ranged from 34.53 to 60.79, indicating varying levels of codon bias, although the overall CUB was weak. CAI values for *MaWRKY* genes ranged from 0.143 to 0.3, suggesting low gene expression levels.

The average content of A3s and T3s was not significantly different from G3s and C3s (P > 0.05) (Fig. 1). However, the combined average content of G3s+C3s diferred significantly from that of A3s+T3s. The average GC content was 55.49%, ranging from 51.05% to 58.95%, indicating a relatively high GC content and a preference for G/C-ending codons in *MaWRKY* genes.

**Fig. 1.**
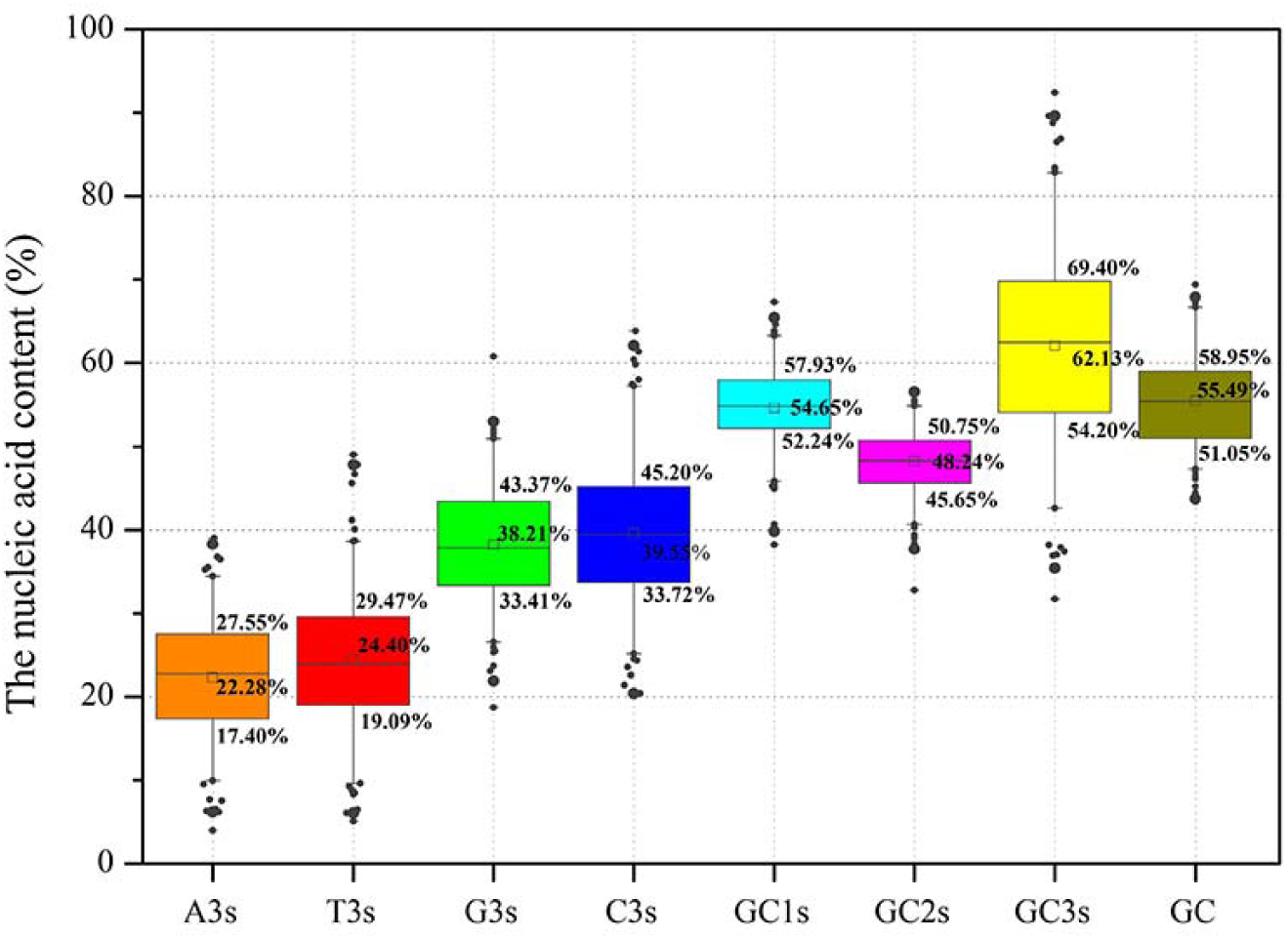
Nucleic acid composition of *MaWRKY* codons. The Y-axis represents the nucleic acid content, while the X-axis corresponds to parameters related to *MaWRKY* codons. These parameters included the total GC content of each coding domain sequence (CDS), the nucleotide composition at the third codon position (A3s, T3s, G3s, C3s), the GC content at each codon position-GC1, GC2, and GC3 (representing the first, second, and third positions, respectively) for each CDS.

Analysis of G/C content at different codon positions showed that GC3s (62.13% on average) were much higher than GC1s (54.65%) and GC2s (48.24%), suggesting preference for G or C at the third codon position. GC3s exhibited greater variability (ranging from 54.20% to 69.40%) compared to GC1s (ranging from 52.24% to 57.93%) and GC2s (ranging from 45.65% to 50.75%). This variability indicates that the first and second codon positions are relatively stable, while the third position shows greater fluctuation, potentially contributing to the observed codon usage bias.

### Codon Usage Parameter Correlation Analysis

The Pearson Correlation coefficients for various codon usage indices were calculated and are presented in Table 1 to assess the relationships between factors influencing codon usage. ENC value showed a significant positive correlation with A3s (r = 0.726, p < 0.05) and T3s (r = 0.690, p < 0.05) but was negatively correlated with G3s (r =-0.616, p < 0.05), C3s (r =-0.703, p < 0.05), GC (r =-0.704, p < 0.05) and GC3s (r =-0.750, p < 0.05). These findings indicate that the nucleotide composition at the third position of synonymous codons strongly influences codon usage bias.

**Table 1.**
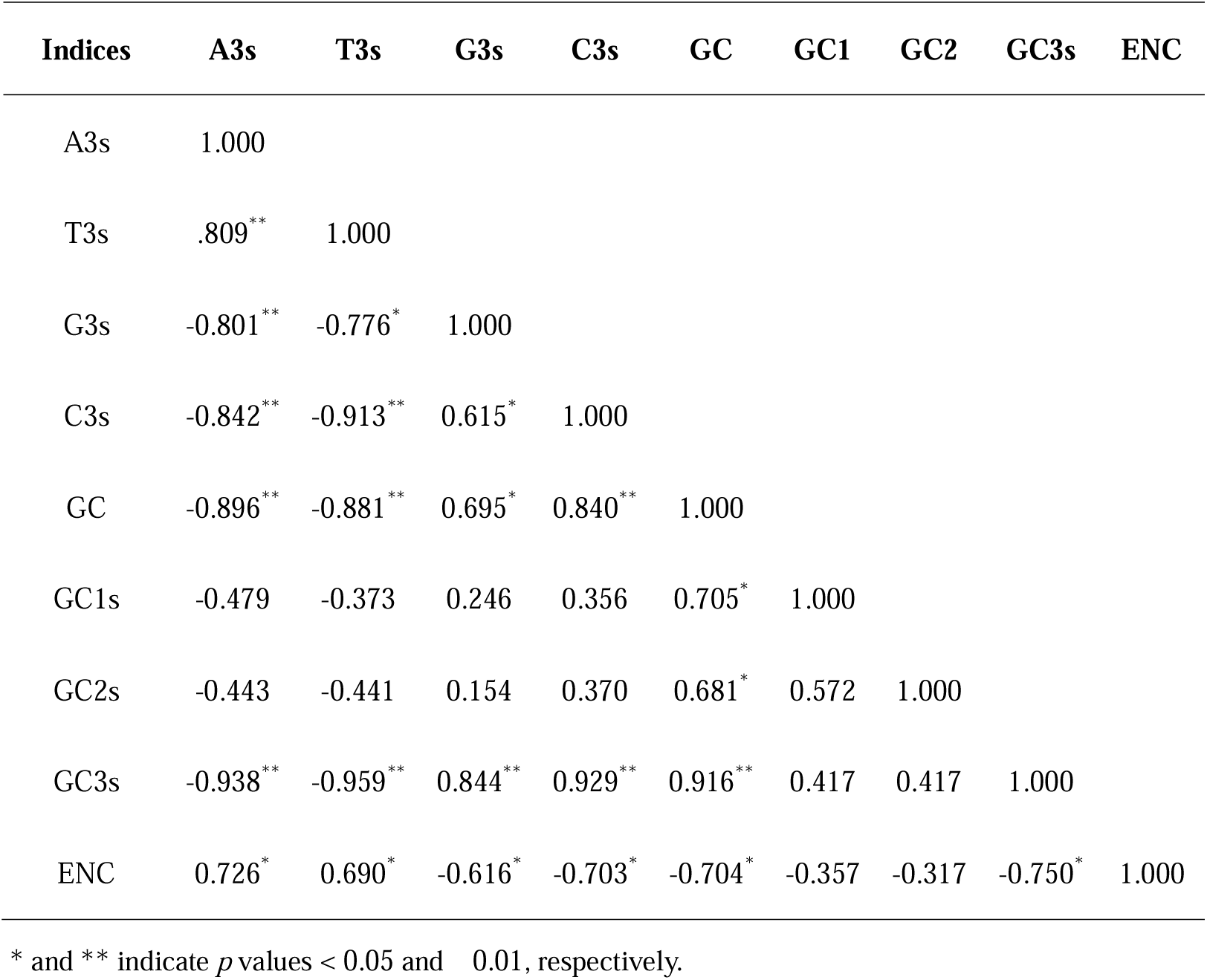
Pearson correlation coefficients among parameters influencing codon bias in *MaWRKY* genes of *Musa acuminata* banana ‘Guijiao 9’. A3s, T3s, G3s and C3s denote the nucleotide content at the third codon position. GC represents the overall GC content of each coding domain sequence, while GC1, GC2, and GC3 refer to the GC content at the first, second, and third codon positions, respectively. ENC indicates Effective Number of Codons.

| Indices | A3s | T3s | G3s | C3s | GC | GC1 | GC2 | GC3s | ENC |
| --- | --- | --- | --- | --- | --- | --- | --- | --- | --- |
| A3s | 1.000 |  |  |  |  |  |  |  |  |
| T3s | .809** | 1.000 |  |  |  |  |  |  |  |
| G3s | -0.801** | -0.776* | 1.000 |  |  |  |  |  |  |
| C3s | -0.842** | -0.913** | 0.615* | 1.000 |  |  |  |  |  |
| GC | -0.896** | -0.881** | 0.695* | 0.840** | 1.000 |  |  |  |  |
| GC1s | -0.479 | -0.373 | 0.246 | 0.356 | 0.705* | 1.000 |  |  |  |
| GC2s | -0.443 | -0.441 | 0.154 | 0.370 | 0.681* | 0.572 | 1.000 |  |  |
| GC3s | -0.938** | -0.959** | 0.844** | 0.929** | 0.916** | 0.417 | 0.417 | 1.000 |  |
| ENC | 0.726* | 0.690* | -0.616* | -0.703* | -0.704* | -0.357 | -0.317 | -0.750* | 1.000 |

\* and \*\* indicate $p$ values $< 0.05$ and $0.01$ , respectively.

Additionally, G3s was positively correlated with C3s (r = 0.615, p < 0.05), GC3s (r = 0.844, p < 0.01) and GC (r = 0.695, p < 0.01). Conversely, A3s exhibited a negative correlation with G3s (r =-0.801, p < 0.01), C3s (r =-0.842, p < 0.01), GC3s (r =-0.938, p < 0.01) and GC(r =-0.938, p < 0.01). Similarly, T3s showed a negative correlation with G3s (r =-0.776, p < 0.05), C3s (r =-0.913, p < 0.01), GC3s (r =-0.959, p < 0.01), and GC (r =-0.881, p < 0.01). These negative correlations suggest that the frequency of A or T at the third codon position has little influence on the codon usage bias of *MaWRKY* genes.

### Principal Component Analysis

Principal Component Analysis (PCA) was performed on the RSCU values of *MaWRKY* genes sequences. The contribution of the first 20 factors to the variance in codon usage bias is shown in Fig. 2A. The first four factors accounted for 58.47% of the total variance, capturing the majority of the differences in codon usage patterns.

**Fig. 2.**
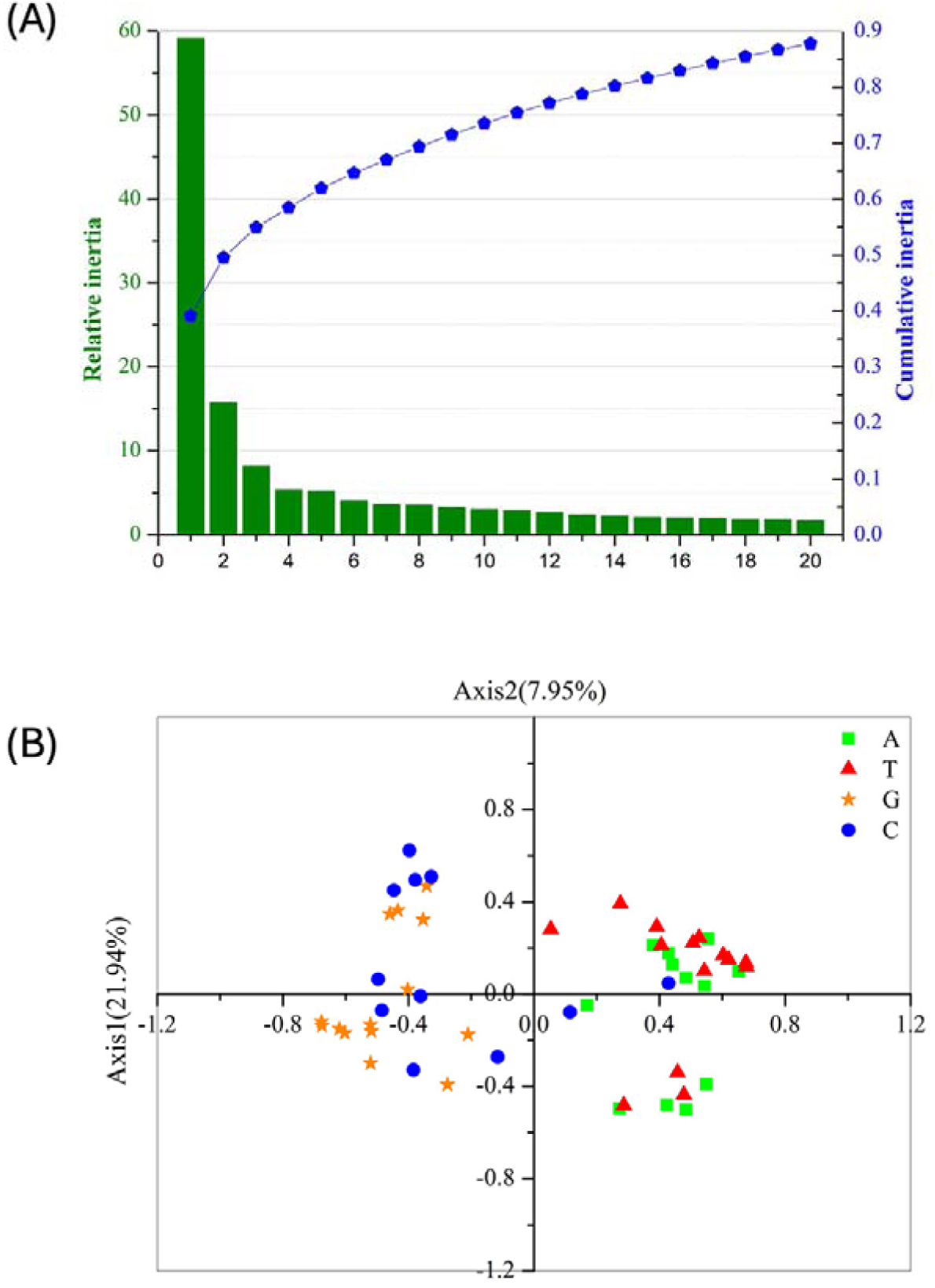
Principal component analysis (PCA) of *MaWRKY* codon usage frequency. (**A)** The relative and cumulative contributions of the first 20 factors to the total variance in PCA; (**B)** PCA plot of codon RSCU values on codons ending with A, T, G, or C. Green squares represent codons ending with A; red triangles indicate those ending with T, orange stars indicate codons ending with G, and blue circles indicate codons ending with C.

PCA analysis of the 59 codons in *MaWRKY* family members, categorised by their endings (A, T, C or G), revealed distinct distribution patterns (Fig. 2B). Codons ending in A and T were clustered on the left side of axis 1, and those ending in G and C were primarily located on the right side. This suggests similar RSCU usage patterns for codons ending in A or T and for those ending in G or C. The distribution of codons ending in A and T was denser and showed partial overlap, whereas codons ending in G and C were more dispersed, with those ending in C being spread across all four quadrants. These findings indicate a natural evolutionary or long-time domestication bias in banana codon usage toward G and C endings.

### Factors affecting codon usage bias in *MaWRKY* genes of *Musa acuminata*

ENC-plot analysis commonly used to assess the effect of mutation pressure on codon usage bias. The ENC values are plotted against GC3 values (Fig. 3A). The standard curve shows that the relationship between ENC and GC3s is shaped primarily by mutation pressure rather than selection. The ENC values for *MaWRKY* family members generally align with the standard curve in the ENC-plot, indicating that mutation pressure is a major factor influencing codon usage bias for these genes (Fig. 3A). However, the ENC values of many *MaWRKY* genes deviate significantly from the standard curve (Fig. 3A), suggesting that mutation pressure is not the sole factor driving codon bias. Other factors, such as natural selection and gene expression, may also contribute to the observed codon bias.

**Fig. 3.**
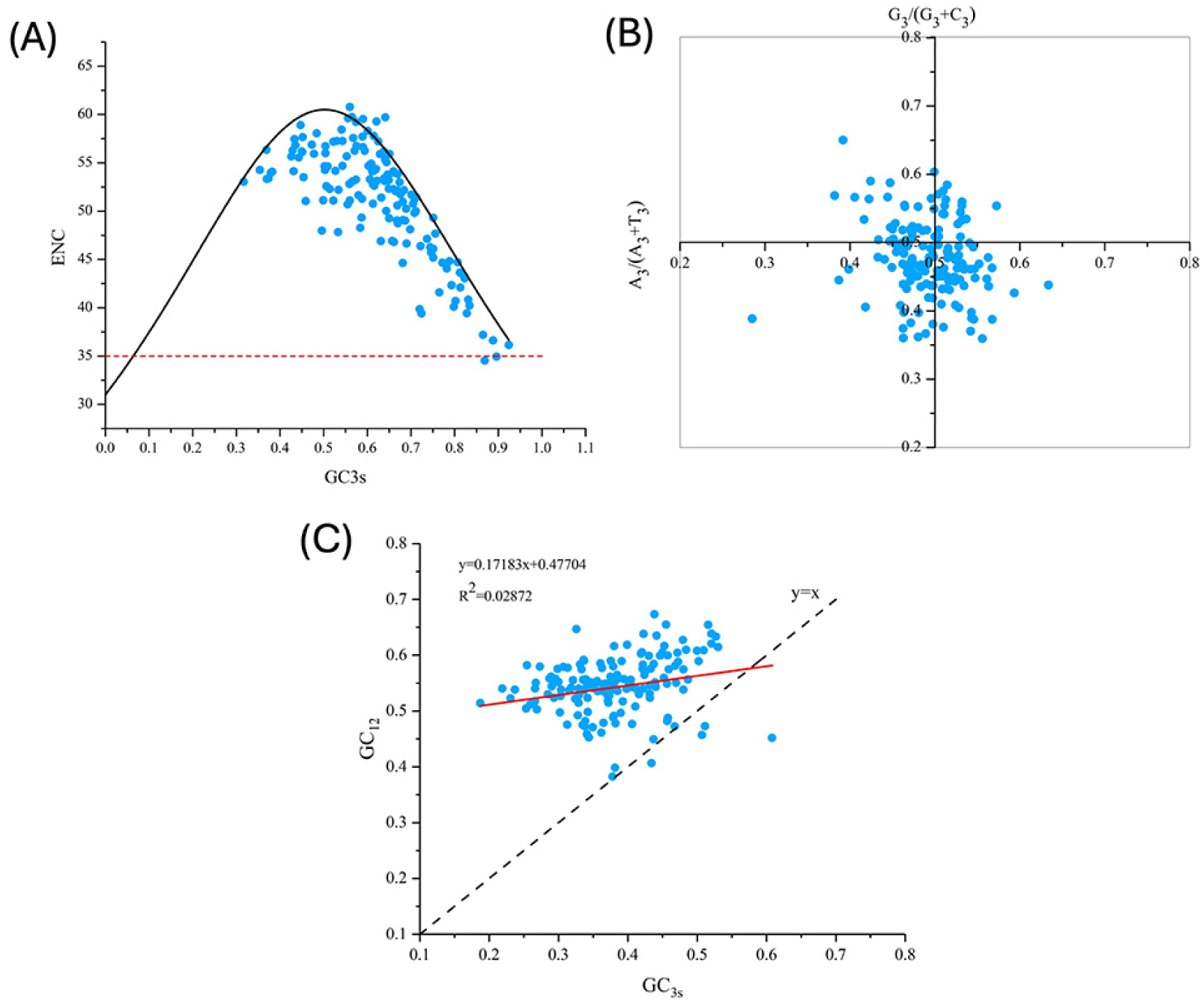
(A)ENC-plot analysis of *MaWRKY* against GC3s. Points on or near the curve indicate bias caused by mutation pressure, while points away from the curve suggest influence from natural selection or other factors. The red-dotted line means the lowest ENC value of *MaWRKY.* **(B)** PR2 plot analysis. The midpoint at 0.5 represents an equal balance between G = C and A = T, indicating no bias between mutation and selection pressure. Deviations from 0.5 suggest that codon bias is primarily influenced by factors other than base mutation of gene-encoding amino acids. **(C)** Neutrality plot analysis of *MaWRKY*. The plot compares GC3 (x-axis) and GC12 (y-axis). An RC value less than 0.5 indicates a stronger influence of natural selection, whereas an RC greater than 0.5 suggests a greater impact of mutation pressure. The red straight line represents the fitted curve.

PR2 analysis was performed to assess the impact of mutation and selection pressure on codon usage by examining whether there was a mutation imbalance between A/T (U) and C/G. In the PR2 plot, A3 / (A3 + T3) is plotted as the ordinate and G3 / (G3 + C3) as the abscissa for *MaWRKY* family members to explore the influence of evolutionary factors (Fig. 3B). The A3 / (A3 + T3) or G3 / (G3 + C3) values for most *MaWRKY* genes deviated from 0.5, suggesting that codon bias was additional influenced by factors other than base mutation of encoding gene amino acids (Fig. 3B). This indicates that additional pressures, such as natural selection, likely played a role in the evolution of *MaWRKY* genes.

To further identify the factors influencing codon usage bias, a neutrality plot analysis was performed by comparing GC3 (abscissa) and GC12 (ordinate) to examine the role of mutation-selection equilibrium in codon usage variation. A linear regression line was plotted with GC3 (abscissa) and GC12 (ordinate) values (Fig. 3C). A regression coefficient (RC) value of less than 0.5 suggests a greater influence of natural selection, while an RC greater than 0.5 indicates a greater impact of mutation pressure. The RC value of 0.1718 indicates that the natural selection had a more significant effect on the codon usage bias of *MaWRKY* gene family in banana than mutation pressure (Fig. 3C).

### Relative synonymous codon usage (RSCU) in *MaWRKY* gene family

To investigate the patterns of synonymous codon usage bias in *MaWRKY* gene family, the RSCU of 59 synonymous codons (excluding the initiation codon, termination codon and tryptophan) was analysed. The results revealed 27 high-frequency codons (RSCU > 1) in *MaWRKY* family members, including AGG, AGA, CUC, CUG, UUG, AGC, UCC, UGG, GUG, GUC, ACC, ACG, GGC, GCC, GCU, CCG, CCA, AUC, UGC, AAG, UUC, AAC, GAG, UAC, CAG, CAC, and GAU, respectively (Fig. 4). Among these, 22 codons ended in G/C, indicating a preference for G or C at the end of the codons in *MaWRKY* members. The codon AGG of Arginine (RSCU = 1.800) exhibited the strongest preference, accounting for 30% of the synonymous codons, while the codon AUC for Isoleucine accounted for 59.0%. These high-frequency codons show preferential usage, which contributed to the deviation of the ENC value of *MaWRKY* genes from an ENC value of 61.

**Fig. 4.**
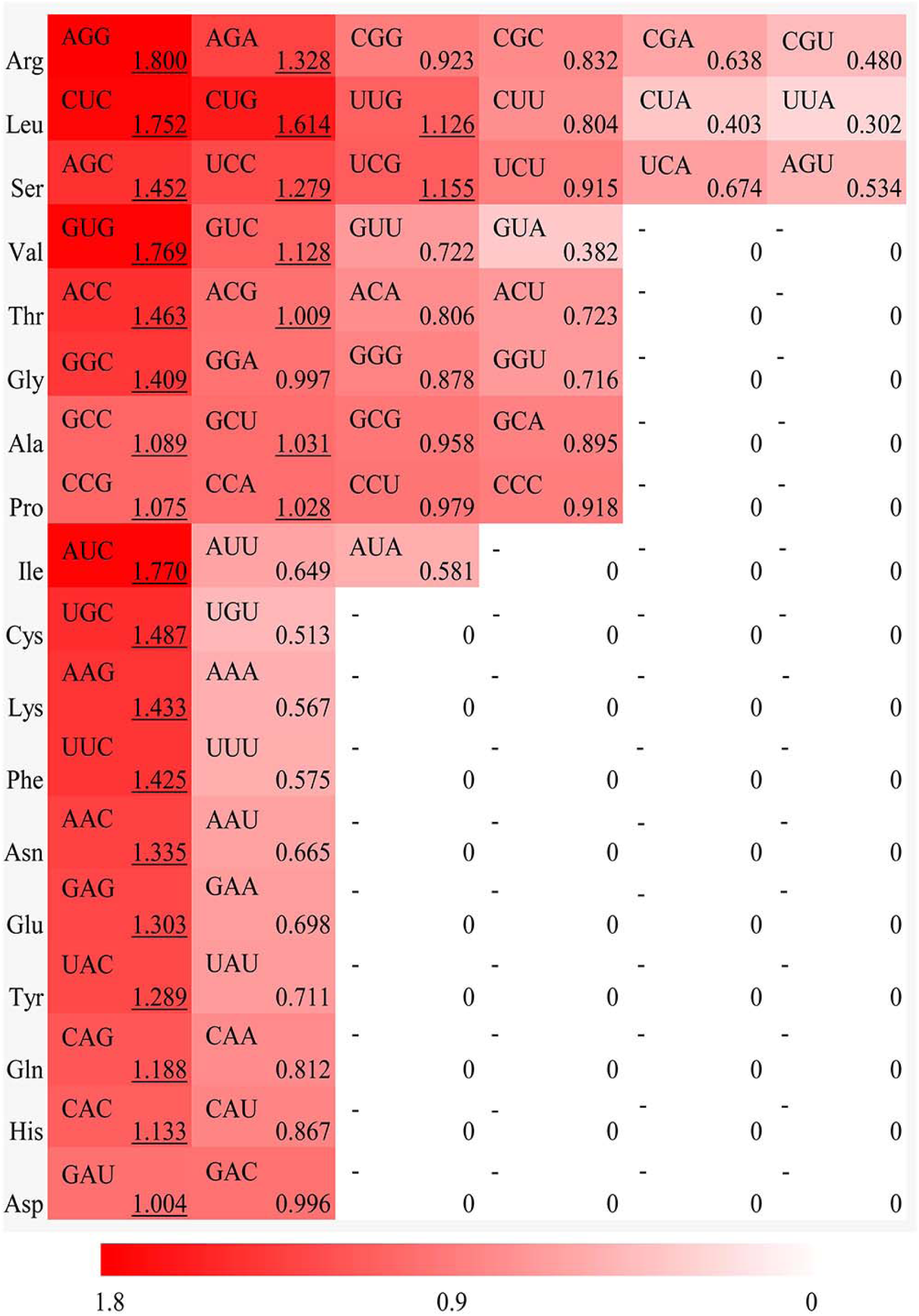
Relative synonymous codon usage (RSCU) analysis for each amino acid in the *WRKY* gene family of *Musa acuminata* banana ‘Guijiao 9’. The red color bar represents the RSCU value, with deeper shades indicating higher RSCU values

### The optimal codons of *MaWRKY* Genes

The optimal codons in *MaWRKY* genes of *M. acuminata* ‘Guijiao 9’ were identified by combining high-frequency codons (RSCU > 1) and highly expressed codon (ΔRSCU ≥ 0.08). The analysis revealed 26 highly expressed codons with ΔRSCU≥ 0.08 (Table 2) and 27 high-frequency codons with RSCU > 1 (Fig. 4). In total, 14 optimal codons were determined for *MaWRKY* genes in *M. acuminata* ‘Guijiao 9’, including CUC, CUG, AUC, GUG, AGC, CCG, ACC, UAC, CAC, CAG, AAC, GAG, UUC and GGC. Notably, all of these optimal codons end with G or C (Table 2).

**Table 2.**
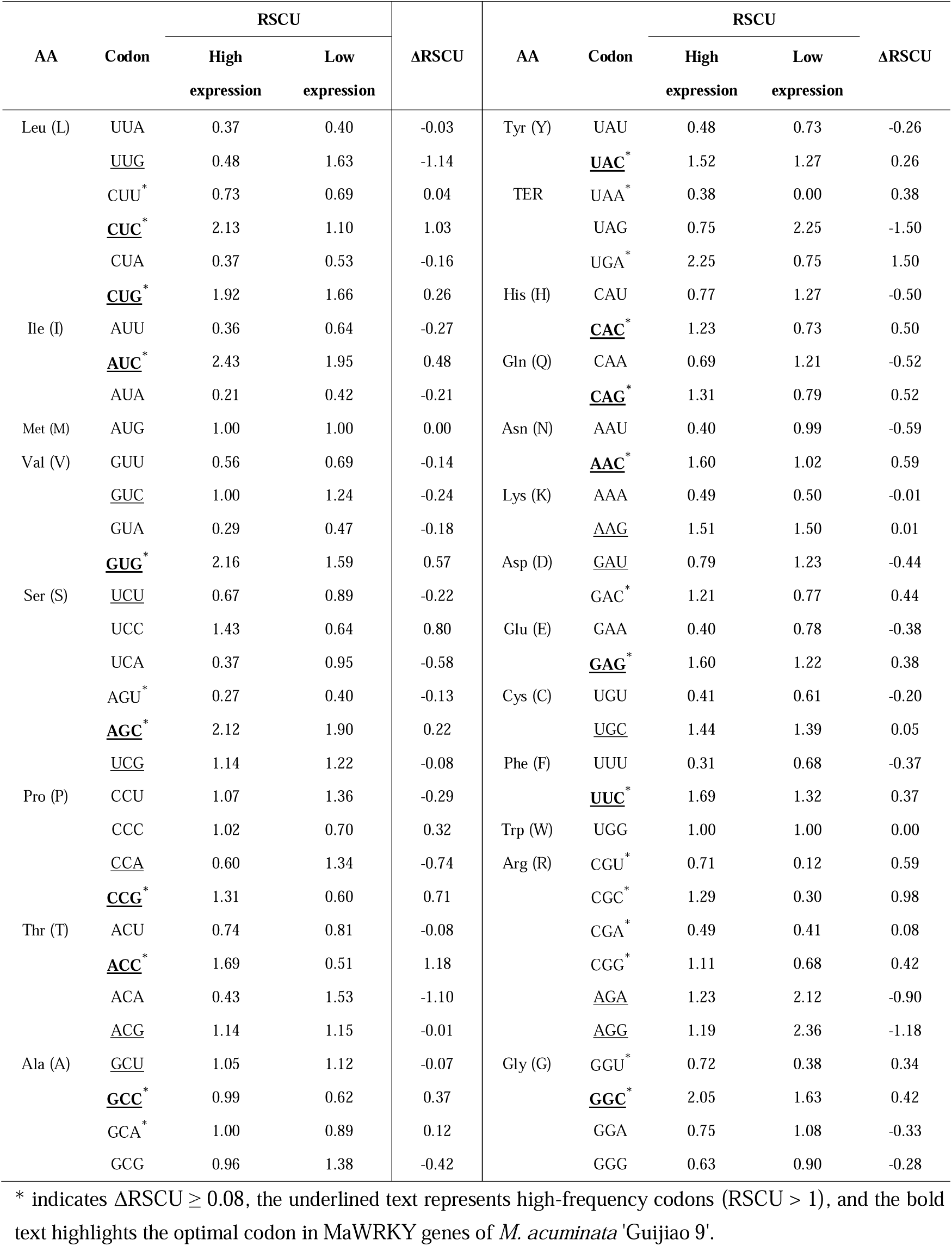
Identification of optimal codons in MaWRKY Genes of *M. acuminata* ‘Guijiao 9’.

### Codon usage patterns of WRKY Family Gene members across different plant species

Using the Euclidean distance, the RSCU values of 25 *WRKY* gene family members across different plant species were clustered with SPSS software v22.0 (Fig. 5). The clustering results of *WRKY* genes in these plant species were consistent with their phylogenetic tree, as shown in Phytozome v12.1 (https://phytozome.jgi.doe.gov/). This finding suggests a correlation between species classification and codon usage patterns. It also indicates that the codon usage patterns of *WRKY* genes in different species align with the evolutionary processes of their respective genomes.

**Fig. 5.**
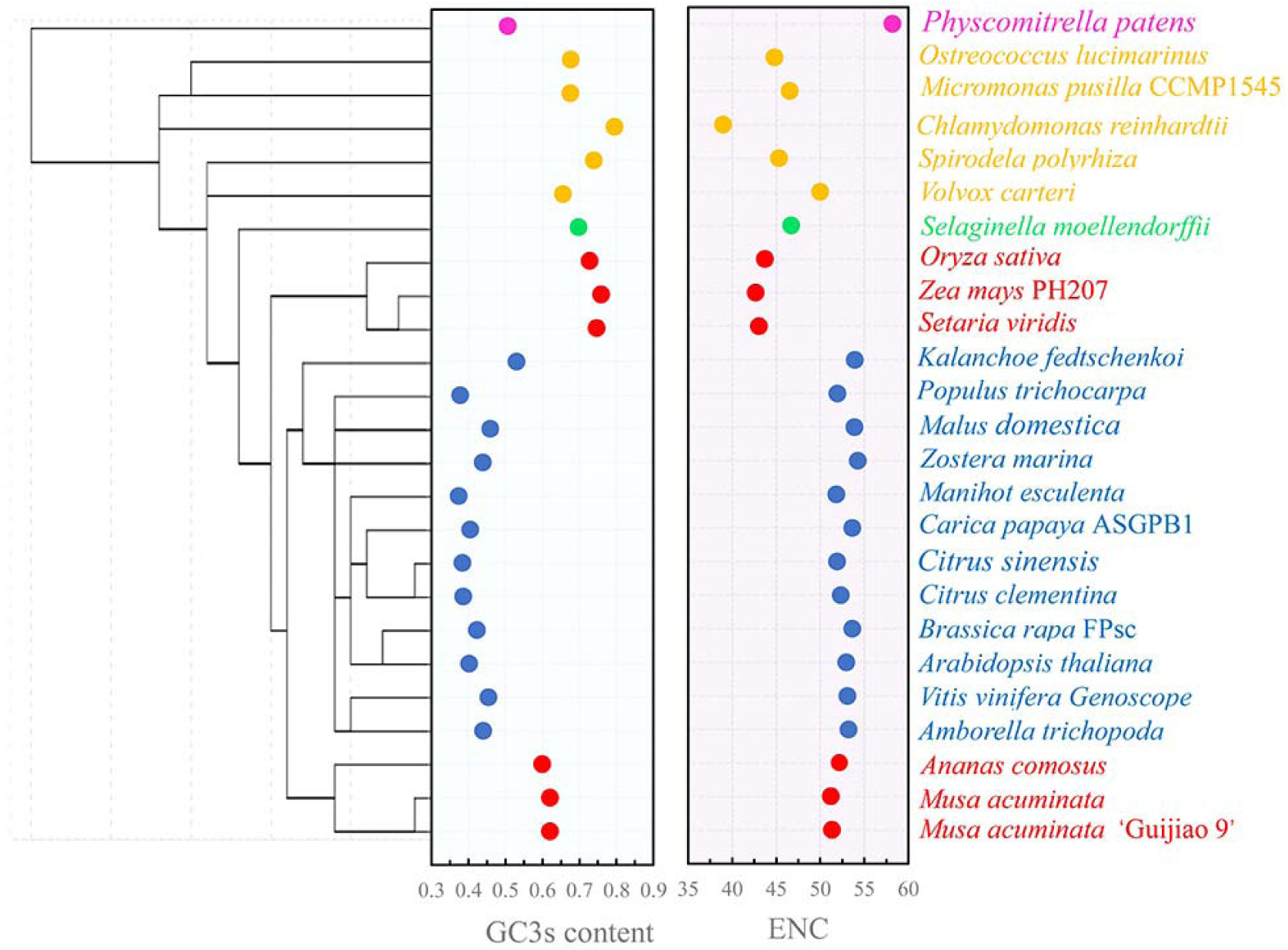
Cluster Analysis of RSCU values of 25 *WRKY* family members across different plant species, along with the mean GC3 and ENC values for each corresponding species. Purple represents Bryophytes, orange represents Algae, green represents Pteridophytes, red represents monocotyledons, and blue represents dicotyledons.

A negative correlation was observed between GC3s content and ENC value when comparing the GC3s and ENC values of WRKY gene family members from different plant species (Fig. 5). The variation in GC3s content and ENC values differed among different plants categories. For example, in monocotyletons, the GC3s content ranged from 60.03% to 75.88%, and the ENC value ranged from 42.67 to 52.20, while in dicotyledons, the GC3s content varied between 37.77% and 52.98%, and the ENC value ranged from 51.85 to 54.26. Plant species that are closely related species showed similar GC3s content and ENC value. ‘Guijiao 9’ and another *Musa acuminata* representative were clustered adjacently due to their similar ENC and GC3 values, as both belong to the monocotyledon group. The ENC value of ‘Guijiao 9’ and *Musa acuminata* is most similar to that of tree species like *Populus trichocarpa*, *Manihot esculenta*, and *Citrus sinensis*, but significantly different from gramineous species such as *Oryza sativa*, *Zea mays* PH207, and *Setaria viridis*.

A heat map analysis of the 59 RSCU values (excluding the initiation codon, termination codon and tryptophan) for 25 plant species revealed that the codons were divided into two groups: those ending with G/C and those ending with A/T. Among the 25 plant species, Algae, Pteridophytes and monocotyledons favoured codons ending in G/C, while dicotyledons preferred those ending in A/T (Fig. 6).

**Fig. 6.**
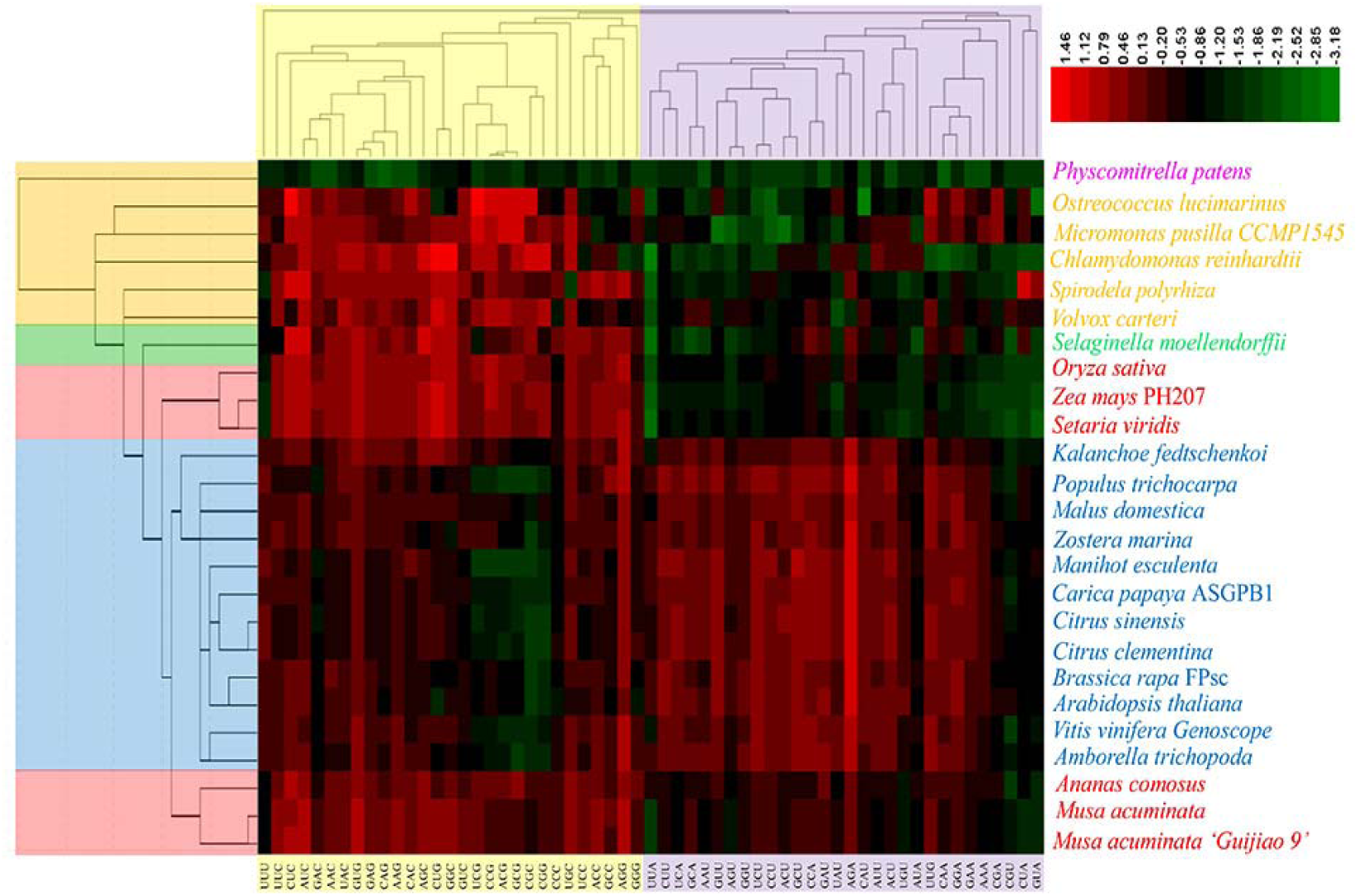
Heatmap analysis of the RSCU values for 25 WRKY family members across different plant species. The color bars represent the RSCU values, with a colour gradient from green to red indicating low to high RSCU values, respectively. The redder the color, the higher the RSCU value, while the greener the color, the lower the RSCU value.

## Discussion

During the process of gene expression, the selective use of codons varies from species to species. Different organisms exhibit different preference for synonymous codons encoding the same amino acid, closely linked to their genetic characteristics. CUB can influence mRNA stability, transcription, protein translation accuracy, and protein folding, thereby fine-tuning gene expression.

In this study, 151 *MaWRKY* genes were identified in *M. acuminata* using bioinformatics tools for codon analysis. GC composition has been shown to influence codon and amino acid usage, with the third base of a codon could directly reflecting codon usage patterns (Chen et al., 2013; Gao et al., 2022). The average GC content and GC3 content of *MaWRKY* codons were 55.49% and 62.13% respectively, indicating a preference for G/C-ended codons over A/U-ended codons in *M. acuminata*. This preference was furher supported by RSCU analysis, which confirmed *MaWRKY* genes favour G/C-ending codons. These results were consistent with the codon usage patterns observed in other monocotyledons, such as in *Musa basjoo*, *Zea mays* and *oryza sativa* (Ma et al., 2015).

The ENC value and CAI value provide insights into CUB and gene expression levels. Higher ENC values suggest lower CUB and gene expression, while higher CAI values indicate better codon adaptiveness and potentially higher gene expression levels. In this study, the *MaWRKY* genes exhibited high ENC values (34.53-60.79) and low CAI values (0.143-0.3), indicating weak CUB and low gene expression levels of these genes in ‘Guijiao 9’. This was consistent with studies showing that most *WRKY* family genes display stress-induced expression patterns (Dong et al., 2003; Kayum et al., 2015). Furthermore, the GC3 content showed greater variability than GC1 and GC2 with codons ending in G/C exhibiting a broad distribution. These findings suggest that nucleotide composition plays a significant role in influencing CUB in *MaWRKY* genes.Mutation pressure and natural selection are recognised as important factors influencing codon usage bias (Sharp et al., 2010; He et al., 2016). In this study, ENC-Plot analysis showed that most ENC values for *MaWRKY* codons did not align with the expected curve, with a few codons exhibiting strong bias near an ENC value of 35. This finding indicates that mutation pressure is not the primary factor driving codon preference. Instead, other factors, such as natural selection and gene expression, likely play significant roles.

PR2 plot analysis further supported this conclusion, as the A3 / (A3 + T3) and G3 / (G3 + C3) values for most *MaWRKY* members deviated from 0.5. This deviation suggests that codon bias in *MaWRKY* genes is influenced more by pressures like natural selection than by mutation. Neutrality plot analysis provided additional evidence, with a regression coefficient of 0.17, highlighting that natural selection exerts a stronger influence on codon bias than mutation pressure.

Base on ENC, PR2 and neutrality plot analyses, the CUB of *MaWRKY* genes in *M. acuminata* is influenced by multiple factors, with natural selection being the dominant force. These findings align with the study by He et al. (2016), which reported that Ginkgo biloba unique genes involved in environment adaptation preferentially using G/C-ending codons, with natural selection as the primary driver of codon usage bias. Interestingly, while monocotyledons like *M. acuminata* favour G/C-ending codons, dicotyledons such as *Helianthus annuus* (Gao et al., 2022) and *Brassica napus* (Li et al., 2013) exhibit a preference for A/T (U)-ending codons. In these species, mutation pressure appears to play a larger role in shaping codon usage bias (Gao et al., 2022; Li et al., 2013). Thus, the codon usage preferences of *WRKY* gene families vary significantly among different species, reflecting distinct evolutionary pressures and adaptations. An optimal codon is defined as one that is over-represented in highly expressed genes and is associated with higher gene expression levels and efficient translation (Zhou et al., 2009). It was generally believed that GC-rich organisms favour codons ending with G or C, while AT-rich organisms prefer codons ending with A or T (Hershberg et al., 2009; Palidwor et al., 2015). In our study, all 14 identified optimal codons of *MaWRKY* genes ended with G or C (Table 2), aligning with the GC-rich composition of *MaWRKY* genes in *M. acuminata.* Similar patterns have been observed in other monocotyledon, such as *Oryza sativa* (Liu et al., 2004) and *Zea mays* (Liu et al., 2010), whose genomes are GC-rich, and whose optimal codons also end with G or C. This finding has practical implications for introducing point mutations and codon modifications to enhance the production of specific proteins in host cells. Codon optimisation based on these preferences can improve gene expression efficiency in biotechnological and agriculture applications.

RSCU values serve as a significant indictor of evolutionary relationships among different organisms and are widely used in studies of biological evolution in plants. Cluster analysis of RSCU values for *WRKY* genes across various plant species revealed that the codon usage patterns of *WRKY* gene families align with the codon usage preferences of their respective genomes during evolution (Fig. 5). *MaWRKY* genes clustered with other monocotyledon species in a single branch, exhibiting a preference for codons ending with G/C (Fig. 6). This suggests that the codon usage patterns of *WRKY* family genes are consistent among related species, reflecting evolutionary.

## Conclusions

In the present study, 151 coding sequences of *WRKY* genes from *M. acuminata* ‘Guijiao 9’ were analysed to investigate codon usage bias and its potential influencing factors. The results revealed that *MaWRKY* genes in *M. acuminata* ‘Guijiao 9’ exhibit a preference for codons ending in G/C. Overall, the codon usage bias was weak, and the expression levels of these genes were low, indicating high variability in synonymous codon usage among *MaWRKY* genes. While mutation pressure played a role, natural selection emerged as the primary factor in shaping the codon usage patterns of *MaWRKY* genes during evolution. The codon usage patterns of *MaWRKY* genes provide valuable insights into their evolutionary dynamics and offer a foundation for identifying and developing suitable gene targets with resistance to biotic stress.

## Materials and Methods

### Screening, Identification and Characterization of Sequence

The complete genome of *M. acuminata* double-haploid ‘Pahang’ was downloaded in FASTA format from the National Centre for Biotechnology Information (NCBI) banana genome database (https://www.ncbi.nlm.nih.gov/datasets/taxonomy/214687/, GenBank assembly accession: GCA_904845865.1), and was used as the reference genome. The Coding Domain Sequences (CDS) of the *WRKY* gene family were retrieved from the transcriptome datat of *Musa acuminata* banana ‘Guijiao 9’ (Additional File 1). Corresponding amino acid and nucleotide sequences were obtained through local BLAST. HMMER 3.3 (http://hmmer.org/download.html) was used to identify sequences containing complete WRKY domain using the reference sequence (PF03106) obtained from PFAM (http://pfam.sanger.ac.uk/). The presence of the WRKY domain in these sequences was further confirmed using NCBI’s CDD tool (https://www.ncbi.nlm.nih.gov/cdd). ORFfinder was then used to identify CDS from the retained sequences (https://www.ncbi.nlm.nih.gov/orffinder/). Ultimately, a total of 151 *M. acuminata WRKY* genes (*MaWRKY)* were identified and included in the subsequent analysis (Additional File 2).

The CDS sequences of the *WRKY* gene family from 24 different species, including Bryophyte, Algae, Pteriophyte, monocotyledon and dicotyledon, were downloaded from the JGI Phytozome12.1 database (https://phytozome.jgi.doe.gov/pz/portal.html) and utilised for Cluster analysis.

### Indices of Codon Usage Bias

Following the screening of *MaWRKY* gene CDS sequences, the key indices of codon usage bias were calculated, including the effective number of codons (ENC), codon adaptation index (CAI), relative synonymous codon usage (RSCU), total GC content of each CDS (GC), and GC content at the first, second and third codon positions (GC1, GC2, GC3). Additionally, the nucleotide composition at synonymous third codon positions (A3s, T3s, G3s, C3s) was analysed. These calculations were performed using the CodonW program (version 1.4.2). Correlations between nucleotide contents were assessed using SPSS statistical software (version 23.0), with values exceeding 0.5 and below-0.5 indicating strong positive and strong negative correlations, respectively.

The Effective Number of Codons (ENC) is a key metric for evaluating the degree of codon usage bias. ENC values range from 20, representing extreme bias where only one codon is used for each amino acid, to 61, indicating no bias where all synonymous codons are equally utilised. Higher ENC values suggest weaker codon usage bias and lower expression levels (Sharp and Li, 1986; Wright, 1990; Zhao et al., 2016). Typically, an ENC value above 40 indicates weak CUB (Wright, 1990). The Codon Adaptation Index (CAI) ranges from 0 to 1, reflecting how well a gene’s codon usage aligns with those of highly expressed genes (Sharp and Li, 1987). Higher CAI values signify better adaptiveness and higher potential expression efficiency. Relative Synonymous Codon Usage (RSCU) measures the observed frequency of a codon relative to its expected frequency, independent of the amino acid composition of gene product (Wang et al., 2016). An RSCU value of 1 signifies no bias, meaning the codon is used as frequently as expected. Values greater than 1 denote a preference for the codon (positive bias), while values below 1 indicate under utilisation (negative bias) (Zhao et al., 2016).

### ENC-plot Analysis

ENC-plot analysis is commonly used to identify factors influencing codon usage bias by plotting ENC values against GC3s values. The standard curve represents the relationship between ENC and GC3s, helping to distinguish whether mutation pressure or natural selection is the dominant factor (Jiang et al., 2008; Liu et al., 2019). When codon usage bias is primarily driven by mutation pressure, the points tend to lie on or just below the standard curve. In contrast, if natural selection and other factors play a larger role, the points will fall below the standard curve (Wright, 1990).

### Parity Rule 2 (PR2) Bias Plot Analysis

A PR2 bias plot is created by plotting the AT bias (A3 / (A3+ T3)) on the y-axis and the GC bias (G3 / (G3 + C3)) on the x-axis. The midpoint at 0.5 indicates an equal balance between G = C and A = T, suggesting no significant bias from either mutation or selection pressure (Deb et al., 2020). If the genes are clustered near the centre, it suggests that the base frequencies are relatively balanced, and the codon bias is mainly affected by mutation pressure. Conversely, If the genes are far from the centre, other factors may be affecting the codon usage bias.

### Neutrality Plot Analysis

A neutrality plot is used to assess the extent to which codon usage bias is influenced by mutation versus selection in organisms, by comparing GC3 (x-axis) and GC12 (y-axis). In this plot, each gene is represented by a point. If the regression coefficient of the plot approaches 1, the points will show a clear pattern, indicating selection plays a significant role in codon usage bias. If the regression coefficient deviates from 1, it suggests that factors other than mutation pressure are also influencing the codon usage bias (Sueoka, 1988).

### Relative Synonymous Codon Usage (RSCU) Cluster Analysis

RSCU cluster analysis was employed to group similar elements. Euclidean distance, a common method for calculating clustering distance, measures the linear correlation between two variables. A smaller clustering distance indicates higher similarity between the clustered genes. The RSCU values of the *MaWRKY* gene family members in ‘Guijiao 9’ were compared with those from 24 other plant species to reveal the evolutionary relationships and characteristics among them.

### Principal Component Analysis (PCA) of RSCU

Principal Component Analysis (PCA) is a multivariate statistical method to examine relationships among multiple variables, and is often applied to analyse trends in synonymous codon usage patterns (Kanaya et al., 1996). In this study, the RSCU values for 59 synonymous codons from *MaWRKY* genes was reduced from 151 dimensions (representing 151 *MaWRKY* genes) to two principal components through dimensionality reduction. Fifty-nine synonymous codons did not include the initiation codon AUG, tryptophan codon UGG, and the termination codons UAA, UAG, and UGA (Zhang et al., 2018).

### Optimal Codon Analysis

The optimal codon plays a significant role in improving the speed and accuracy of translation (Duret et al., 1999). To predict the optimal codon for *MaWRKY*, the ΔRSCU method was applied (Qiu et al., 2011). The *MaWRKY* gene sequences were sorted based on their ENC values. The top 10% and the bottom 10% of the genes, based on ENC value, were selected to create two new databases (high expression and low expression, respectively), and the RSCU values for the codons were calculated. A codon was considered a high-frequency codon if its RSCU value was greater than 1, while a codon was regarded as a high-expression codon if the ΔRSCU value was equal to or greater than 0.08. A codon is defined as optimal if it met both criteria: RSCU> 1 and ΔRSCU≥ 0.08.

### Statistical analysis

The CDS sequences of the *MaWRKY* were analysed using CodonW (http://codonw.sourceforge.net/) and EMBOSS online software (http://www.bioinformatics.nl/emboss-explorer/). Codon-related parameters were statistically analysed using Microsoft Office Excel 2016 and MeV software version 4.9.0 (www.tm4.org/mev.html) was used to construct the heat map. ENC-plot analysis, Neutrality plot, and Box diagram were generated using Origin 8.0 and HemI 1.0 (http://hemi.biocuckoo.org/down.php) software. Clustering analysis and principal component analysis(PCA)of RSCU values for *WRKY* genes in different plant species were performed by Statistical Product and Service Solutions (SPSS) version 22.0.

## Abbreviations

CUB: Codon usage bias
ENC: Effective number of codons
CAI: Codon adaptation index
RSCU: Relative synonymous codon usage
PCA: Principal component analysis
PR2: Parity Rule 2
GC1, GC2, GC3: The GC content at the first, second and third position, respectively
A3s, T3s, G3s, C3s: The content of each nucleotide of the codon at synonymous third positions

## Supplementary Information

**Additional file 1.** The RNA-Seq data of *Musa acuminata* banana ‘Guijiao 9’

**Additional file 2.** The composition indices values of codon usage in MaWRKY genes

## Authors’contributions

Conceptualization, Funding acquisition, Investigation, Methodology, Project administration, Resources, Supervision, Writing - original draft, and Writing - review & editing: J.S.; Data Curation, Formal Analysis, Investigation, Methodology, Visualization, Writing - review & editing: J.Z.; Formal Analysis, Investigation, Supervision, and Writing - review & editing: Jinzhong Z; Supervision, and Writing - review & editing: B.J.F.; Formal Analysis, Investigation, Supervision, Writing - original draft, and Writing - review & editing: A.C. All authors have read and approved the final manuscript.

## Availability of data and materials

The genome datasets generated and analysed in the study are available in NCBI, https://www.ncbi.nlm.nih.gov/datasets/taxonomy/214687/, and supplementary files.

## Declarations

## Ethics approval and consent to participate

Not applicable.

## Consent for publication

Not applicable.

